# Precision Fermentation of Recombinant Myofibrillar Proteins for Future Foods

**DOI:** 10.64898/2026.04.20.719284

**Authors:** James Dolgin, Cornelia H. Barrett, Maxwell J. Nakatsuji, Juan Aguilera-Moreno, David L. Kaplan

## Abstract

Myofibrillar proteins, namely actin and myosin, are responsible for many of the textural attributes of animal-based meat. Precision fermentation (recombinant production of food ingredients) represents an underexplored approach to producing these proteins without the unsustainable practice of animal agriculture. We show that through the solubility-enhancing SUMO peptide tag and precipitation-based purification, we can produce actin via recombinant DNA methods at titers of 326 mg/L *E. coli* culture. We also show expression and precipitation of a recombinant fragment of the myosin tail, leading to 572 mg/L culture. For both proteins, yields are improved compared to prior studies, without the need for low-yielding laborious purification columns, with final purities of 69-73%. These recombinant actin and myosin proteins showed macro- and microscopic fibrous features similar to meat. When combined with plant-based proteins, chewiness, hardness, and Young’s modulus were improved towards that of animal-based meat. Preliminary cost analyses suggest a less expensive process for producing myofibrillar proteins compared to established methods. Our results reveal a novel scalable approach to making meat-like foods and ingredients through precision fermentation.

## INTRODUCTION

Precision fermentation, the process of generating animal-based ingredients in microbial cell hosts, is a novel biotechnological method for animal-free protein production.^1^ While conventional animal agriculture is associated with numerous negative externalities, including land use inefficiency,^2^ greenhouse gas emissions,^3^ zoonotic disease,^4^ and animal suffering,^5^ precision fermentation represents a promising approach to produce animal-based ingredients more sustainably, efficiently, and ethically. Precision fermentation has been used to create various animal-based proteins for animal-free food products such as casein (cheese), β-lactoglobulin (milk), ovalbumin (egg), and hemoglobin (meat blood).^6^ However, the use of precision fermentation for producing myofibrillar proteins, namely actin and myosin, remains underexplored. Myosin and actin are the primary constituent myofibrillar proteins in meat, forming sarcomeres which give rise to the hierarchical structure of myofibers and whole-muscle tissue. These myofibrillar proteins are largely responsible for meat’s textural, and gelling properties.^7^ Myosin forms gels upon heating through crosslinking of head and tail regions,^8^ giving meat its chewiness and hardness when cooked. Fibrillar actin (F-actin) contributes viscoelastic properties such as stiffness and binds volatile compounds to enhance meat’s organoleptic qualities.^9,10^ Myosin and actin also both bind to food-relevant proteins and carbohydrates, and can thus serve as healthy protein-rich binders and texturizing additives.^9,11–14^

Producing these proteins through precision fermentation could create novel animal-free foods with enhanced taste and texture, accelerating adoption of alternative non-animal proteins.

While many meat-alternative foods currently exist, many fall short of replicating their animal-based counterparts. Plant-based meat analogues, which aim to mimic the taste, texture, and appearance of meat, have become widely popularized in the past decade. Still, excitement for these meat-like substitutes has waned significantly in recent years, primarily due to shortcomings in sensory attributes, concerns with overprocessing, and dissimilarity to animal-based meat.^15^ Plant-based proteins are limited to a globular morphology, which does not mimic the texture and mouthfeel of fibrillar meat proteins.^16^ This explains much of the sensory gap between plant-based and livestock-based meat. To mimic the texturizing properties of meat, plant-based foods require chemical additives and processing which compromise taste and nutritional value.^17^ Additives such as methylcellulose and carrageenan provide no nutritional benefits and have even been cited for having serious gastrointestinal health risks.^18^ Other approaches to creating realistic meat alternatives include “cultivated meat,” which involves the *in vitro* growth and differentiation of animal cells. However, this approach suffers from numerous biological and engineering challenges.^19,20^ It also still requires the use of livestock for initial cell biopsy, making it non-vegan. Through precision fermentation of myofibrillar proteins, we can generate the proteins responsible for meat’s taste and texture at lower cost, faster turnaround, and without the need for cell biopsy.

While actin and myosin have been expressed recombinantly, current yields and purification processes are insufficient for the food industry. Protein yield, the final amount of target protein produced per volume of culture, is balanced with purity, the ratio of target protein to total protein. Attempts to produce and purify recombinant actin, whether from bacterial or yeast host organisms, have yielded a maximum of 10 mg/L culture.^21–25^ This stems from actin’s low solubility and high loss to affinity columns leading to low recovery. Recovery is defined as the ratio of target protein present at the end versus the start of a purification process. Myosin has also proven difficult to express and purify due primarily to the protein’s size (∼223 kDa) and complexity of head region folding.^26^ Some have solved this issue by expressing only fragments of the tail region of myosin,^27–29^ the region responsible for myosin’s mechanical and gelling properties.^30^ However, after purification steps, this still leads to yields of 10 mg/L or less.

Column chromatography and ultracentrifugation, the usual purification techniques for these proteins, lead to high protein loss and low yields, and are very expensive to scale.^31,32^ To produce recombinant myofibrillar proteins for food applications, more efficient purification methods are needed.

Here, we explore production improvements to recombinant actin and myosin and their use as precision-fermented food proteins. For protein production we use *E. coli* strain BL21 and closely related strain BLR, which are well-characterized, non-pathogenic strains^33^ with FDA-approval for food ingredient production.^34,35^ We show how soluble yields of recombinant actin can be improved through fusion tags (SUMO), and a more scalable purification process using ammonium sulfate precipitation. For recombinant myosin, we demonstrate that an optimized 99 kDa fragment of the myosin tail produces high yields and forms stable gels. We demonstrate that these proteins form fibrillar meat-like structures, with actin polymerizing to form F-actin bundles, and myosin forming fibers upon freeze-drying. Finally, we show that these myofibrillar proteins are able to improve the textural properties of soy protein isolate, demonstrating their potential as binders and texturizing agents in plant-based foods. These results show recombinant myofibrillar proteins as a potential new breakthrough in alternative proteins, creating taste and texture-relevant animal proteins through economical and efficient processes without the need for animal slaughter or animal cell culture.

## EXPERIMENTAL SECTION

### Results

#### Yield and purification process for recombinant actin is improved using SUMO tag and ammonium sulfate precipitation

To improve soluble yield of recombinant bovine actin from *E. coli,* an N-terminal small ubiquitin-like modifier (SUMO) peptide tag was employed, leading to a 5.8-fold increase of soluble actin compared to non-SUMOylated actin (Fig. 1a). The SUMO tag, which increases solubility through favorable protein folding,^36^ can easily be cleaved off by Ulp1 peptidase, which scarlessly cleaves the SUMO tag through recognition of its tertiary structure. Scarless cleavage is important for processing precision fermented proteins, as non-native amino acid sequences lead to increased allergenicity risks.

**Figure 1:**
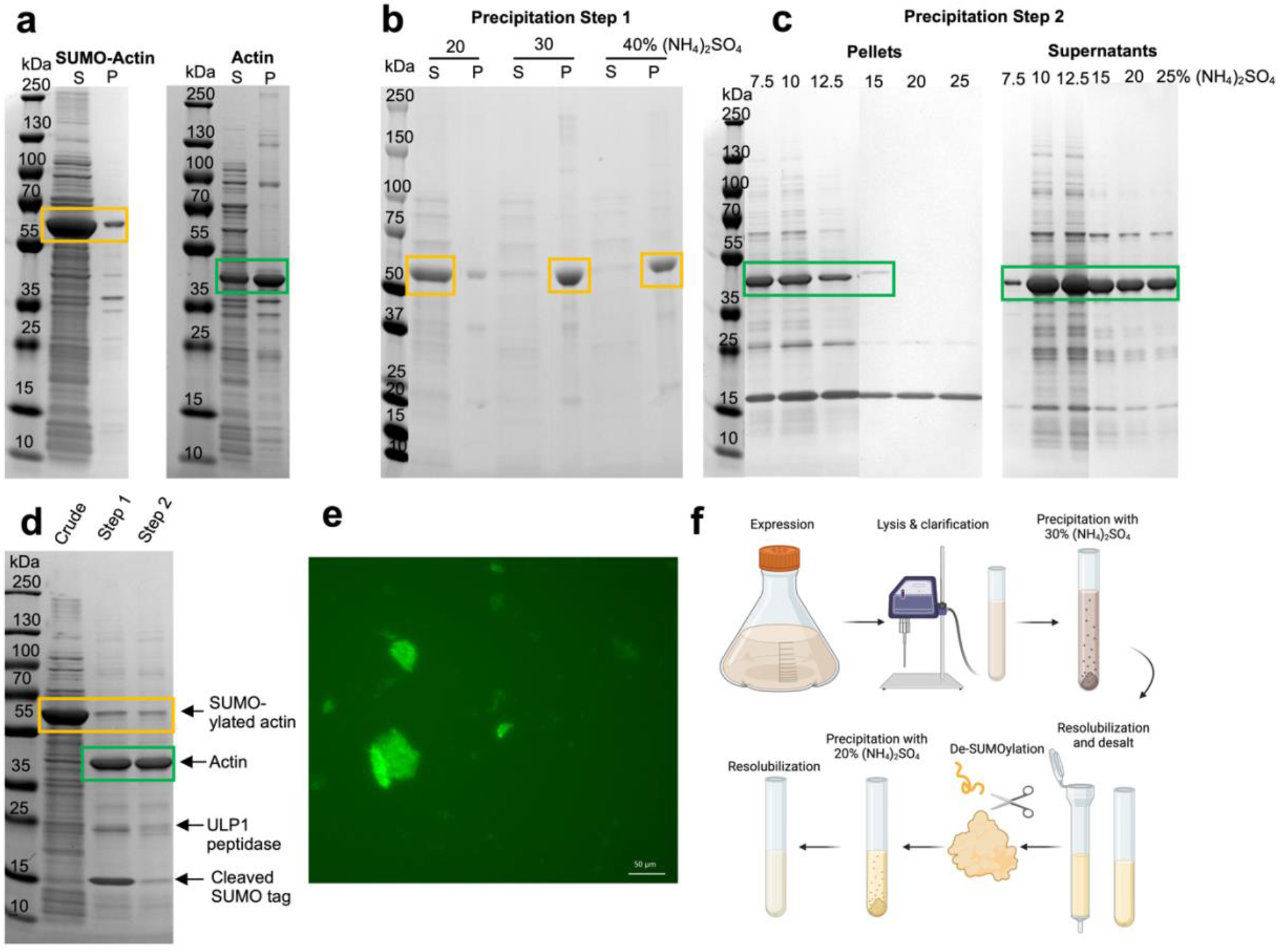
Production of recombinant actin using SUMO tag. **a)** SDS-PAGE showing soluble supernatant (S) and insoluble pellet (P) after lysis and centrifugal separation of SUMO-Actin compared to neat actin. **b)** SDS-PAGE showing various concentrations of saturated ammonium sulfate used to pellet SUMO-actin from crude supernatant, showing supernatant (S) and pellet (P) after precipitation and centrifugation. **c)** SDS-PAGE showing various concentrations of saturated ammonium sulfate used to pellet actin after enzymatic cleaving of SUMO tag, with pellet and supernatant separated to show gradual increase in pelleting with ammonium sulfate increase. **d)** SDS-PAGE showing complete purification process of recombinant actin, starting with unpurified SUMO-Actin (“Crude”), actin after 30% ammonium sulfate precipitation, solubilization, and cleaving with ULP1 peptidase (“Step 1”), and actin after subsequent 20% ammonium sulfate precipitation to remove ULP1 and SUMO tag (“Step 2”). **e)** Filamentous actin after polymerization and staining with green fluorescent phalloidin (SB = 50μm). **f)** Overview of complete purification process for recombinant actin (created with biorender.com). Yellow boxes indicate SUMOylated actin; green boxes indicated non-SUMOylated actin.

Ammonium sulfate ((NH_4_)_2_SO_4_ ) precipitation was used to salt out our protein of interest. Ammonium sulfate is low cost, and is a generally recognized as safe (GRAS) substance widely used in the food industry.^37^ Addition of saturated (NH_4_)_2_SO_4_ at 30% volume was determined to be optimal for initial precipitation from crude protein lysate (Precipitation Step 1), as shown by SDS-PAGE as near-complete removal of SUMO-Actin from supernatant (Fig. 1b). After cleaving the SUMO tag, we successfully removed much of the residual enzyme and tag through optimizing saturated (NH_4_)_2_SO_4_ to 20% volume addition (Fig. 1c, Precipitation Step 2). The protein content of the crude versus purified fractions can be seen in Fig. 1d, showing a final protein purity of 72.5%, in line with that of established precision-fermented protein ingredients.^32,38^ Final yields are 326 mg actin per 1 liter culture with recovery of 45.9%, significantly higher than established yields and recoveries.^21,31^ We confirm successful production of actin through polymerization and staining with phalloidin (Fig. 1e), which selectively targets filamentous actin. We show that the use of a SUMO tag improves yields and processing compared to actin expressed without such tag (Table 1), with recovery and purity higher in SUMO-actin compared to neat actin. Neat actin also required more (NH_4_)_2_SO_4_ (50% vs 30%) to be precipitated (Fig. S1). The full purification process can be seen in Fig. 1e. Our results ultimately show that a combination of SUMO-tagging and (NH_4_)_2_SO_4_ precipitation of recombinant actin result in a translatable process for alternative protein applications, with significant improvements to yield and recovery compared to conventional column-based procedures.

**Table 1:**
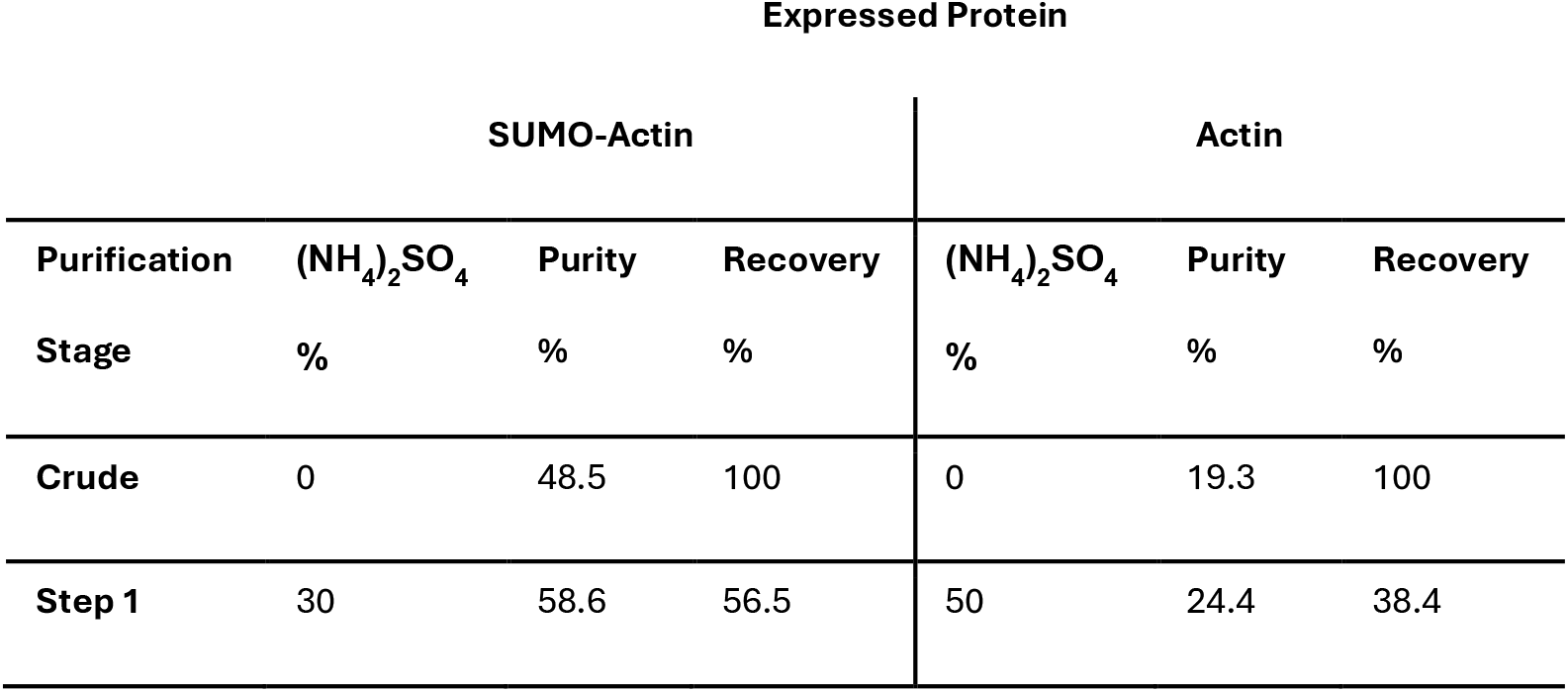
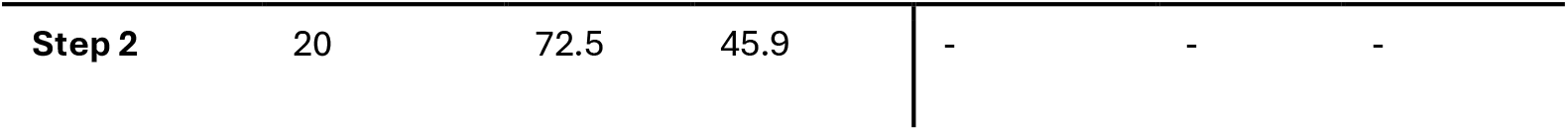
Purity and recovery of SUMO-actin or actin at stages of protein purification, and ammonium sulfate ((NH_4_)_2_SO_4_ ) concentration used at each step.

#### A 99 kDa myosin tail fragment allows for simple expression and purification using buffer exchange

To determine which fragment of the myosin tail region to express for downstream applications, we expressed C-terminal fragments of sizes 64, 99, and 131 kDa (Fig. 2a) bovine myosin in *E. coli*. We desired stable expression and heat-inducible gelling of the expressed fragment to show behavior akin to meat protein. The 64 kDa region was initially selected due to previously documented success expressing a similar sized fragment of rabbit myosin in *E. coli*,^29^ and gelling of similarly sized myosin fragments in fish surimi products.^39^ Though expression of the 64 kDa region was successful (Fig. 2b), the expressed protein did not gel upon heating as indicated by the inverted test tube technique (Fig. 2c). This may be due to different behaviors of myosin in fish and mammalian organisms. A larger 131 kDa myosin tail fragment, representing the entire light meromyosin region, was expressed. This led to inefficient expression (Fig. 2b), likely arising from internal translation due to the many methionine residues present in this fragment being misinterpreted as start codons. To solve this problem, we selected an alternative downstream start site with a high translation rate using RBS designer^40^ to express the 99 kDa myosin tail fragment. This fragment expressed well according to SDS PAGE and formed a stable gel upon heating (Fig. 2b & 2c). This indicated 99 kDa myosin could be produced recombinantly with *E. coli* and mimic meat properties for alternative proteins.

**Figure 2:**
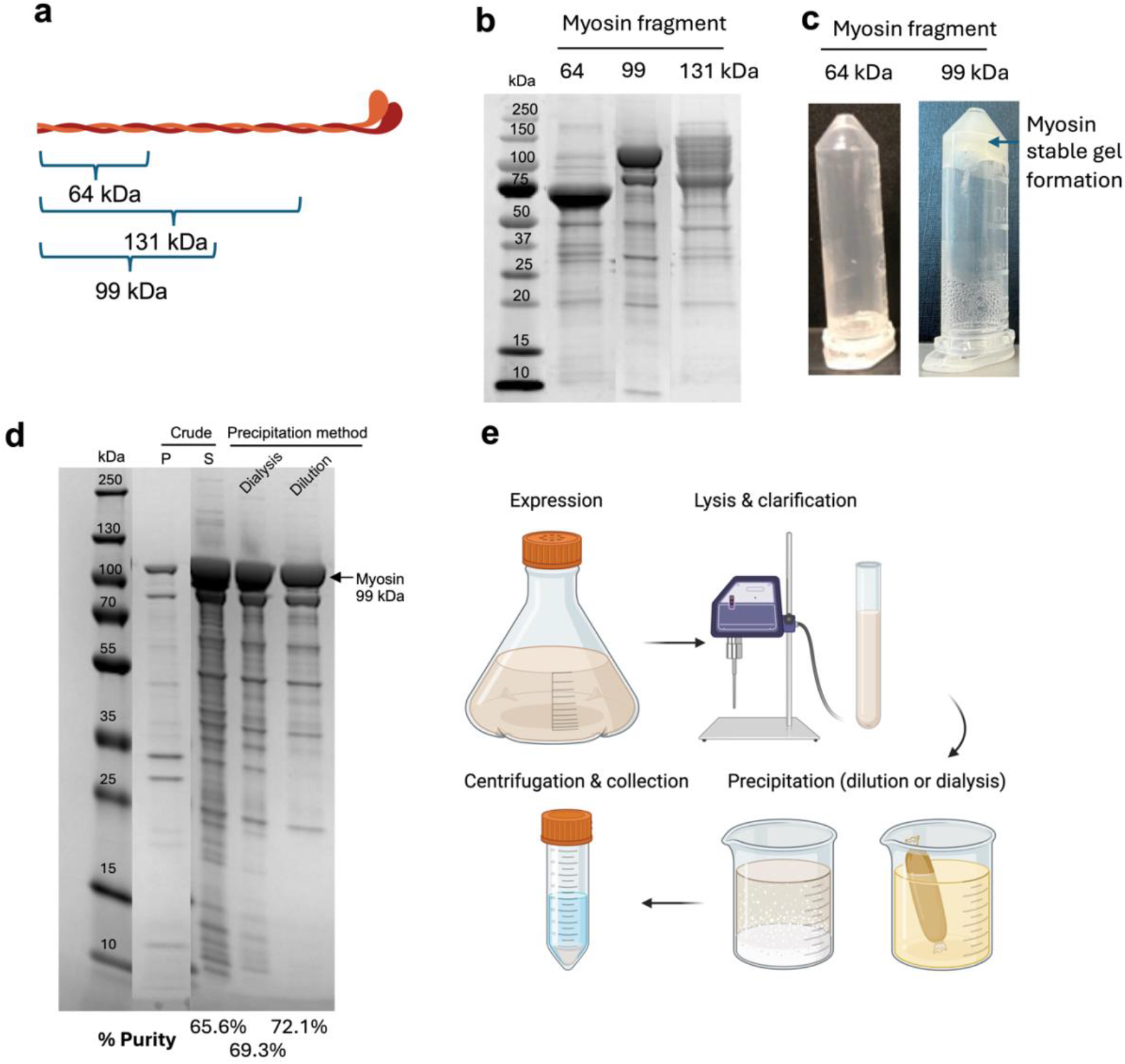
Production of recombinant myosin tail fragment. **a)** Schema representing approximate size and location of expression sites in myosin protein. **b)** SDS-PAGE showing unpurified supernatants from expression attempts of myosin fragments. **c)** Inversion assay to test gel-forming ability of 64 and 99-kDa myosin fragment upon heating, showing stable gel formation of 99-kDa fragment but not 64-kDa fragment. **d)** SDS-PAGE of 99-kDa myosin fragment after lysis and centrifugal separation into pellet (P) and crude supernatant (S), followed by collection of protein through precipitation using dialysis and dilution methods, with associated purities indicated. **e)** Overview of complete purification process for recombinant myosin 99-kDa fragment. Figures a & e created with biorender.com.

To concentrate and purify the 99 kDa myosin fragment from crude supernatant, we took advantage of myosin’s insolubility in low salt. After lysis, myosin was precipitated from supernatant with either dialysis or dilution to create a low-salt environment. After centrifugation and supernatant removal, this led to myosin purities of 69.3% (dialysis method) or 72.3% (dilution method) (Fig. 2d). While dilution is a simpler and more scalable process, dialysis led to ∼2.9 fold more concentrated myosin. Depending on final application, one process may be preferred to the other. Final calculated yields were 572 mg/L culture with 90.4% recovery from crude supernatant. The full process for recombinant myosin production can be seen in Fig. 2e.

#### Hierarchal fibrous structures achieved by recombinant myofibrillar proteins

An important feature of muscle tissue is its hierarchal structure which gives meat its unique texture. Micron-scale myofibrils organize into muscle fibers tens of microns wide, and these structure sizes have significant impacts on meat quality.^41^ We show that our recombinantly produced myofibrillar proteins form similarly hierarchal fibrillar structures. When polymerized, fibrillar actin forms bundles on the tens of microns scale, with 45.3 ± 22.2μm diameter (Fig. 3a). Individual fibers, when imaged via SEM, show <10μm diameter (Fig. 3b), in line with reported results of *in vitro* polymerized actin.^42^ We found that upon lyophilization, expressed 99 kDa myosin formed a fibrous structure that would be expected from whole-cut meat products (Fig. 3c). These fibers comprised a wide range of sizes, from sub-micron to >10 micron, and organize into porous structures similar to freeze-dried muscle tissue^43^ (Fig. 3d). Myosin formed these structures upon drying while actin did not, perhaps due to self-assembly of coiled-coil tertiary structures unique to myosin. Actin and myosin formed fibers of similar diameters, but myosin fibers had larger diameter range (4.4 ± 1.8μm for actin, 4.2 ± 4.1μm for myosin; Fig. 3e).

**Figure 3:**
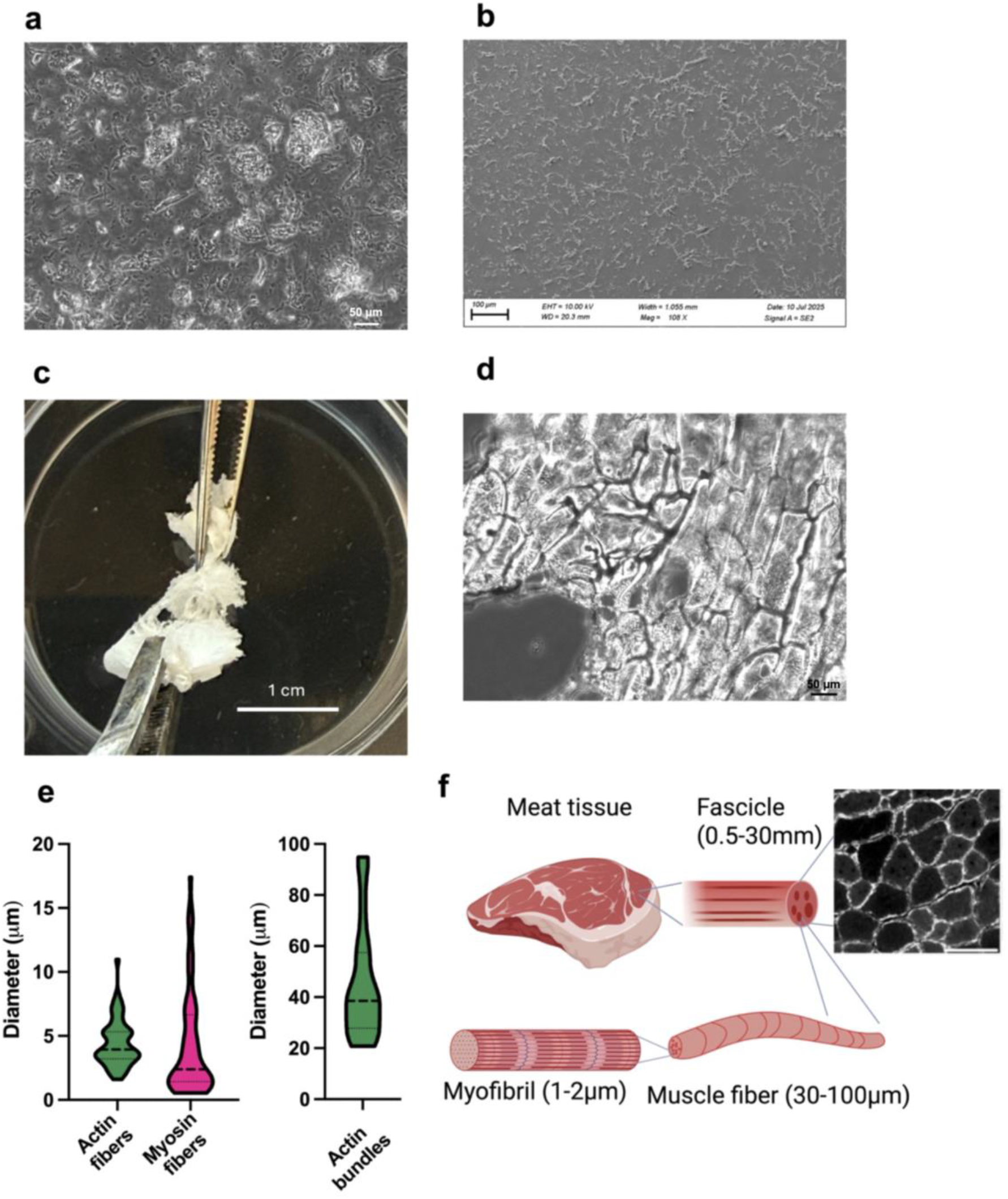
Morphology of recombinant myofibrillar proteins. **a)** Brightfield image of filamentous actin in solution after polymerization (SB = 50μm). **b)** Scanning electron microscopy image of desiccated actin fibers to resolve individual fibers (SB = 100μm). **c)** Digital photograph of lyophilized 99-kDa myosin sample after dialysis purification, showing fibrous structure. **d)** Brightfield image of lyophilized 99-kDa myosin (SB = 50μm). **e)** Image-based quantification of actin and myosin fiber diameters (n=60), and diameter of actin bundles (n=30). **f)** Representation of muscle tissue hierarchical structure (created with biorender.com), with histological cross-section of muscle fibers shown. Reproduced with permission from ref.^43^ (Copyright 2021, https://creativecommons.org/licenses/by/4.0/; SB = 250 μm)

Overall, these recombinant myofibrillar proteins cover a range of fiber sizes for mimicking the fiber sizes and morphology of animal-based meat (Fig. 3f), from the sub-micron scale to tens of microns, allowing for the formulation of a broad range of potential alternative protein products.

#### Recombinant myofibrillar proteins modulate texture of plant-based protein

Current plant-based meat alternatives lack the textural properties of animal-based meat.^44^ We show that recombinant myofibrillar proteins can improve these properties in plant-based protein. Using texture profile analysis (TPA), we evaluated the effect of adding varying amounts of our recombinantly produced fibrillar actin and 99 kDa myosin to soy protein isolate (SPI). SPI and myofibrillar proteins were homogenized and heated and samples were subjected to a double compression test to acquire results (Fig. S2). We also conducted this test on ground sirloin beef for comparison. Results show that hardness of samples was significantly improved even by addition of 0.2% weight myosin (Fig. 4a). At 1% weight actin and myosin, hardness was improved by several-fold, bringing hardness closer to that of sirloin beef. Cohesiveness, related to a food’s consistency, was improved by adding 0.3% actin but decreased by adding 0.3% myosin (Fig. 4b). A similar trend was observed in measurements of resilience (Fig. 4c), related to a food’s mechanical recovery after compression. These trends could be due to rigid short actin fibers resisting microcrack formation in the SPI matrix under compression, as can be seen in fiber-reinforced materials.^45^ Springiness, a food’s ability to maintain “bounce” after multiple compressions, was decreased by smaller myosin additions but unchanged by actin addition (Fig. 4d). This could be due to a dominance of myosin’s viscous properties at lower concentrations. Chewiness, the product of hardness, cohesiveness, and springiness, is an important metric in evaluating texture as it relates to the overall tenderness of food. Addition of actin and myosin led to significant improvements to chewiness, bringing samples within the range of animal-based beef (Fig. 4e). Young’s modulus (stiffness) was also improved at just 0.1% addition of actin, and improved further at higher concentrations of actin and myosin (Fig. 4f).

**Figure 4:**
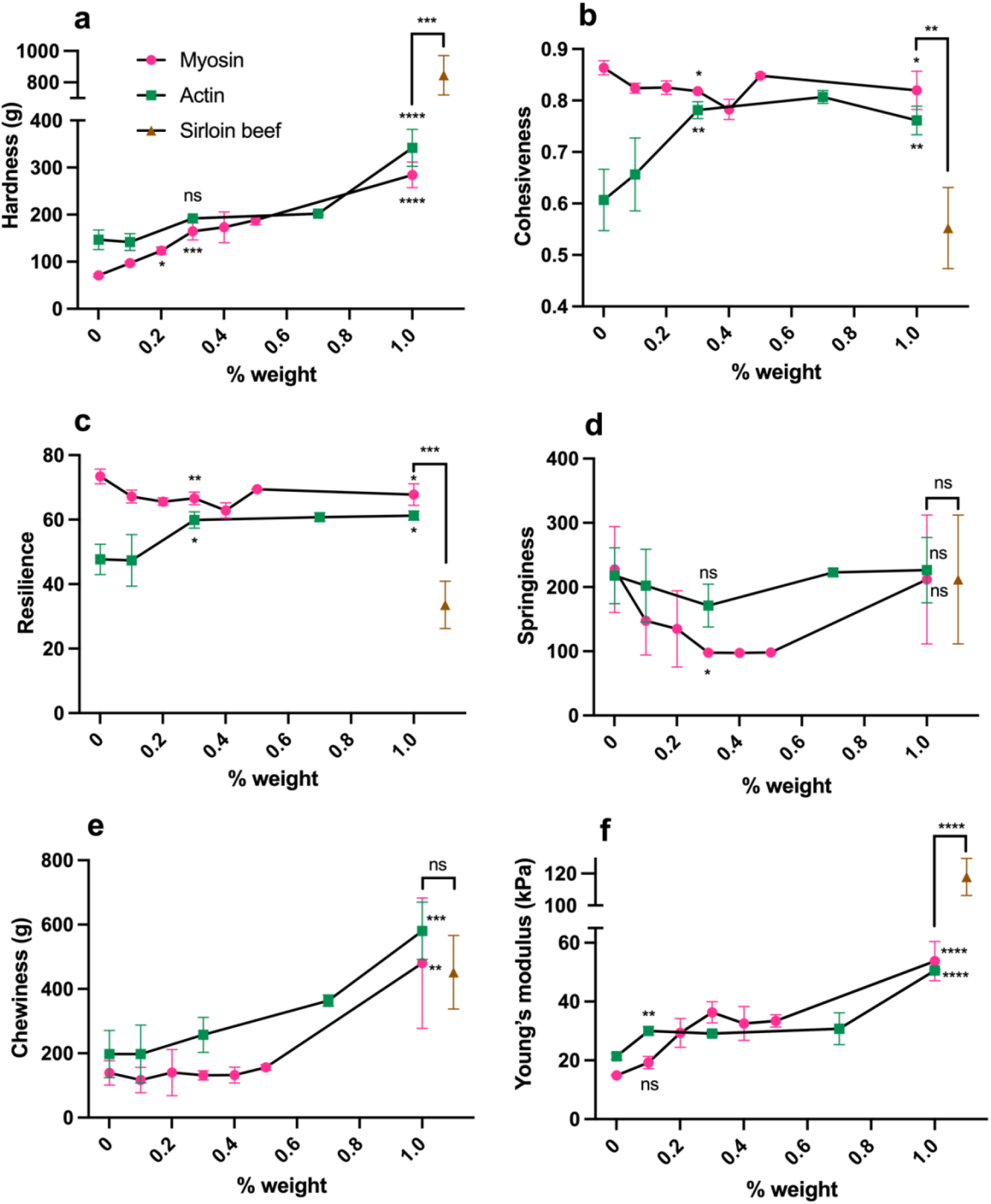
Textural properties of soy protein isolate (SPI) cooked with varying concentrations of recombinant fibrillar actin or myosin 99-kDa fragment, compared to cooked ground sirloin beef. Graphs show **a)** hardness, **b)** cohesiveness, **c)** resilience, **d)** springiness, **e)** chewiness, **f)** Young’s modulus. % weight represents target protein mass/total sample mass. Adjacent asterisks represent comparison to 0% weight control. All presented data is mean values±SD. Statistical analysis by one-way ANOVA with Dunnett’s correction for multiple comparisons, with p-value cutoffs of >0.1234 (ns), 0.0332 (*), 0.002 (**), 0.0002 (***), <0.0001 (****).

Overall, recombinant actin and myosin significantly improved textural properties of plant-based protein, showing they can be used as a helpful ingredient in the formulation of meat alternatives. Myofibrillar proteins interact with plant-based protein via sulfhydryl bonds,^9,11–14^ and contribute to heat-based gelling and stiffness properties.^7,10^ This likely led to the observed changes to hardness, cohesiveness, springiness, chewiness, resilience, and stiffness of SPI, with concentration-dependent impacts to properties. Myofibrillar proteins brought properties like hardness, chewiness, and Young’s modulus closer to those of cooked ground beef. Chewiness of SPI was made comparable to ground beef by using 1% recombinant actin or myosin. Textural properties of meats vary widely depending on species and processing,^44^ so the tunability of texture with recombinant myofibrillar proteins offers valuable customization of food products.

#### Recombinant production saves cost and time compared to cultivated meat

Growing meat-relevant proteins using cell culture, the process known as cultivated meat, is a widely explored approach to creating animal-free alternative proteins.^20,46^ Here, we compared this strategy of producing myofibrillar proteins to precision fermentation. After culturing and differentiating immortalized bovine satellite cells (iBSCs) on adherent plates, we extracted whole-cell protein and determined myofibrillar protein content. We found there to be 57.5 mg myosin and 32.5 mg actin per 175 cm^2^ flask (Fig. S3). Immunostaining of myosin heavy chain in iBSCs validated successful differentiation. To achieve the equivalent yield of 1L precision fermented myosin, we estimate 10 flasks (1,750 cm^2^) of iBSCs must be grown and differentiated over a period of 8 days (4 days of proliferation, 4 days of differentiation), compared to the 3 days needed for precision fermentation (Fig 5a).

**Figure 5:**
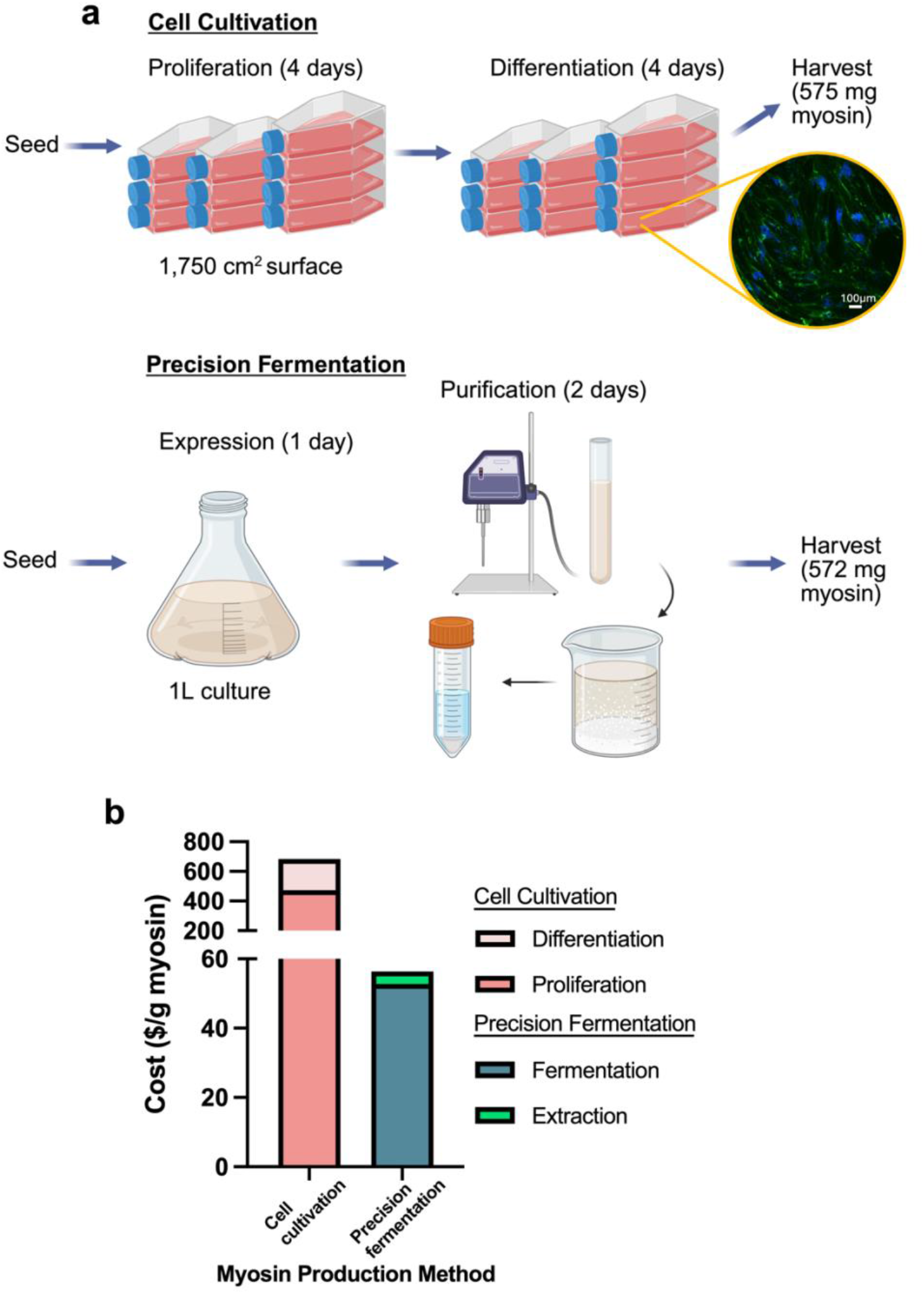
Comparison of myofibrillar protein production using cultivated meat (cell cultivation) versus precision fermentation. **a)** Illustration of lab-scale process needed to generate similar myosin yield using either cell cultivation of immortalized bovine satellite cells (iBSCs) or precision fermentation. (created with biorender.com) Pop-out shows myosin heavy chain immunostaining of differentiated iBSCs (green = myosin heavy chain; blue = nuclei; SB =100μm). **b)** Raw materials cost estimate of lab-scale production of 1g myosin using either cell cultivation or precision fermentation.

We also determined cell cultivation to be far more expensive than precision fermentation for myofibrillar protein production. We estimated lab-scale raw materials cost for production of 1g myosin to be $56.38 via precision fermentation and $683.35 via cell cultivation (Fig. 5b).

Production costs for cell culture were driven primarily by fetal bovine serum (FBS) and differentiation factors (Table S1). Most production costs for precision fermentation were due to fermentation broth and induction reagent (Table S2). These costs would decrease significantly at scale, especially with decreased feedstock costs, enhanced strain engineering, and optimization of protein output.^6^ Cost estimates did not include estimates for utilities, labor, water, consumable materials, sterilization, and equipment, though these costs are likely to be higher for cultivated meat which is highly susceptible to contamination, requires highly specialized bioreactors, and has high energy and water usage demands.^19,47^

## Discussion

Myofibrillar proteins, namely actin and myosin, contribute significantly to the textual and organoleptic qualities of meat.^7,9^ Through precision fermentation, these proteins can be produced cheaply and efficiently without the use of animal livestock to create sustainable alternative protein foods. Current approaches to recombinant production of actin and myosin rely on column chromatography and ultracentrifugation, processes that are challenging to scale and which result in low yields and high production losses.^21–29^ We show that soluble yield of recombinant actin can be improved with a SUMO peptide tag, and this actin can be simply purified using two steps of ammonium sulfate precipitation. We also show that the functional tail-portion of myosin can be expressed effectively in its 99-kDa form and purified with buffer exchange. Our criteria for purity was >65%, as determined by standards set by the Food and Drug Administration (FDA) vis-à-vis the safety evaluation of Impossible Foods’ precision-fermented heme protein.^48^ There is no established minimum purity for approval of precision fermented products, leaving some leeway for future directions in lower purity and, in turn, higher yield. Recombinant actin purification, for example, may be simplified to only one step of ammonium sulfate precipitation which would improve yield. To further simplify processing, low-cost plant-derived enzymes and their associated recognition sites can be utilized to remove the SUMO tag instead of the ULP1 peptidase. In-house production of ULP1 was straightforward but would represent additional costs at scale. Further optimization of myosin tail fragment expression is also worth investigation, possibly through a more granular library screening approach to optimize expression and mechanical properties, as has been previously explored with elastin-like peptides.^49^ Overall, our improved processes for myosin and actin precision fermentation represent significant steps toward using these proteins in an alternative protein context.

We then evaluated the textural impact of recombinant myofibrillar proteins, analyzing their morphological characteristics as well as their influence on mechanical properties of meat alternatives. Sensory perception of food depends heavily on particle size^50^ and shape.^51^ Actin fibers, formed upon polymerization of recombinant actin, ranged from 2-10μm, forming higher-order bundles 20-100μm in diameter. Myosin fibers covered a range of sizes 1-20μm wide with fibrous structures visible to the naked eye. These fibers also formed a honeycomb-like structure similar to that of animal muscle tissue. Animal muscle tissue exhibits a hierarchal structure of 1-2μm myofibrils making up the 30-100μm diameter muscle fibers found in meat,^52,53^ similar to the fiber sizes formed by these recombinant proteins. These micron-scale structures cannot be recapitulated in the extrusion processes conventionally used for making fibrous textured plant-based meat, as these processes form fibers of much larger diameters.^54^ Recombinant myofibrillar proteins’ ability to mimic the micron-scale morphology of animal-based meat is significant considering the micron-scale sensitivity of human taste perception. Using specific combinations of myosin and actin, particle size distributions can be further tuned, impacting perceptions of tenderness, juiciness, and water holding capacity.^50^ These proteins can be used as enhanced-mouthfeel ingredients or standalone food products with the fibrous quality of animal-based meat, though further studies should explore the textural and sensory qualities of such standalone food products compared to meat.

Sensory perception also depends heavily on mechanical properties, which we evaluated using texture profile analysis (TPA). TPA determines food’s mechanosensory properties using a double compression test, calculating relevant characteristics from stress and strain.^20^ To show the potential use of recombinant myofibrillar proteins as texturizing ingredients, we cooked these proteins together with soy protein isolate (SPI), a common plant-based meat substitute, at varying concentrations. We found that hardness, chewiness, and Young’s modulus, which plant-based meat analogues lack compared to animal-based meat,^44^ were improved with myofibrillar protein addition, bringing these properties closer to those of ground beef. Cohesiveness, resilience, and springiness could be differentially modulated depending on concentration and choice of myosin or actin. Control groups for myosin and actin differed in some properties due to slight differences in buffer compositions. Texturizing binders such as polysaccharide gums and cellulose derivatives are often included in plant-based meat analogues at 1-3% composition,^55,56^ but these ingredients raise concerns of inflammation, indigestion, and overall lack of nutritional value.^18,57^ The recombinant proteins used in this study showed greater impacts to textural properties using lower concentrations compared to conventional binders.^55,56^ This potency could be due to the heat-based gelling properties of myofibrillar proteins as well as their binding strength to plant-based proteins through sulfhydryl bonds.^9,11–14^ They also offer greater potential nutritional benefits than conventional binders, as they are complete protein sources which should metabolize as conventional meat proteins. Overall, recombinant myofibrillar proteins have great potential as texture-enhancing ingredients for plant-based foods. They improve important textural properties that plant-based foods otherwise lack and can tune texture to create custom alternative protein products. Myosin or actin can be selected and used according to specific textural or particle size needs in final products. Actin, for example, can be used to create stiffer product at lower concentration, while myosin can impart meat-like fibrousness and substantial improvements to hardness. Combining myosin and actin together may synergistically improve mechanical properties, as the two proteins are strong binding partners. Follow-up studies should explore this approach for creating potent myofibrillar food ingredients. Translation from laboratory to table remains a challenge for precision-fermented proteins due to cost and scalability obstacles. We estimated lab-scale production costs of 1g recombinant myosin to be $56.38. Actin costs will be greater due to needs of recombinant ULP1 peptidase, desalt columns, and ATP buffer components. These costs would significantly decrease at scale and with further optimization. Techno-economic analyses show that through improvements to expression titer, cost reduction of carbon source, and process optimization, recombinant protein costs can be brought to $37/kg using *E. coli*, and $10/kg using fungal systems,^58^ which would make costs competitive with certain polysaccharides like carrageenan and pectin.^59^ Recombinant myofibrillar proteins may also be used as ingredients in cultivated meat. We estimated cultivated meat to cost $683.35 per gram myosin, though these costs can also decrease significantly with scale and process optimization.^60^ Cultivated meat production is time-consuming and final products lack the texture of animal-based meat.^20^ Recombinant myofibrillar proteins can improve sensory qualities in cultivated meat and replace lengthy differentiation processes, creating new opportunities for precision fermentation-cultivated meat hybrid products.^46^

In conclusion, precision fermented myofibrillar proteins represent a unique opportunity to produce animal-like protein-rich foods without conventional animal agriculture. We demonstrate approaches to significantly improve the production and purification process of these recombinant proteins through peptide tags, fragment expression optimization, and precipitation techniques. We then show how these proteins can improve textural properties by forming unique morphologies and mechanical property improvements of plant-based proteins. This is shown to be a lower-cost approach to myofibrillar protein production than cultivated meat.

## Methods

### Plasmid construction

All plasmids were codon-optimized for *E. coli* expression and synthesized using ThermoFisher Scientific GeneArt service unless otherwise noted. Gibson assemblies were carried out using New England Biolabs (NEB, Ipswich, MA) Gibson Assembly Master Mix (NEB #E2611) and manufacturer protocols. Ligations were carried out using NEB KLD enzyme kit (M0554) and manufacturer protocols. Primers were designed and PCR conditions determined using NEBuilder. PCR reactions were carried out using NEB Q5 High-Fidelity DNA Polymerase (NEB #M0491) and dNTPs (NEB #N0447). Plasmids were expanded using 5-alpha competent *E. coli* (NEB #C2987) in LB Broth (Sigma Aldrich, St. Luois, MO, #3522) with kanamycin 50μg/mL (Sigma Aldrich #K4000) as antibiotic, and purified using Macherey-Nagel (Düren, Germany) NucleoSpin Plamid Mini kit (#740588). Plasmid insertions were verified using Azenta (Burlington, MA) Genewiz Sanger sequencing service using T7 promoter and terminator as forward and reverse primers, respectively.

Actin sequence was optimized to express bovine alpha skeletal muscle actin (UniProt ID: P68138). Actin DNA fragment was cloned downstream of SUMO sequence on vector pCIOX, a gift from Andrea Mattevi (Addgene plasmid # 51300), using primers in Table S3 followed by Gibson Assembly and direct transformation into 5-alpha *E. coli*. SUMO sequence was deleted, using primers in Table S3, and re-ligated for non-SUMOylated expression from the same plasmid.

Sequence for 64-kDa myosin was optimized to express bovine myosin 1 (UniProt ID: Q9BE40) starting from lysine residue 1389. This sequence was cloned into pCIOX vector in place of the actin sequence without SUMO using primers in Table S3 with Gibson Assembly. Myosin construct for 131-kDa fragment was created by synthesizing codon-optimized sequence of residues 811-1388 using plasmid synthesis service from GenScript (Piscataway, NJ), and Gibson Assembly to previously created 64-kDa myosin sequence using PCR with primers in Table S3. Finally, 99-kDa myosin plasmid was synthesized by deleting residues 811-1089 and re-ligating.

The full sequence for codon-optimized actin and myosin fragments used in this study can be found in Table S4. The associated plasmid construct for SUMO-Actin and 99-kDa myosin can be found in Figure S4.

### Actin protein production

Plasmids containing sequence for bovine actin or SUMO actin were transformed into *E. coli* BLR (Sigma Aldrich # 69053) and transformed bacteria were kept on kanamycin-containing agar-LB broth plates at 4°C or in 30% glycerol (Sigma Aldrich G6279) for long-term storage at - 80°C. Auto-induction media (Boca Scientific, Dedham, MA, # GCM19, 55.6 g/L), supplemented with 4mL/L glycerol and 50μg/mL kanamycin, was inoculated with overnight culture to optical density 600nm (OD600) of 0.1. After overnight expression, cells were spun down with centrifugation (4,000xG 10 min.) and pellets frozen at -80°C, then resuspended in lysis buffer to OD 30. Lysis buffer contained 50mM Tris-hydrochloride (Tris-HCl, pH 8), 0.2mM calcium chloride (CaCl_2_), 0.2mM adenosine-triphosphate (ATP from pH 8 stock), 1mM Tris(2-carboxyethyl)phosphine (TCEP), and 0.6% sarkosyl. Cells were lysed using a Branson (Brookfield, CT) SFX 550 sonifier using a 3s on/9s off cycle at 25% amplitude for 15 minutes total. Protein was precipitated using saturated ammonium sulfate by dropwise addition while stirring on ice for 20 minutes. Precipitated protein was spun down with centrifugation (16,000 xG 15 min.) and resuspended in G-Actin (globular actin) buffer, containing 5mM Tris-HCl (pH 8), 0.2mM CaCl_2_, 0.2mM ATP, 0.1 mM TCEP. Protein was stirred at 4°C until fully dissolved, then desalted using a Zeba Spin Column (10mL, 7k kDa cutoff, Fisher Scientific) equilibrated with G-Actin buffer to remove residual ammonium sulfate. SUMO tag, if present, was cleaved using 32 μg/mL ULP1 peptidase overnight at 4°C. Protein was re-precipitated with ammonium sulfate, resolubilized, de-salted, and polymerized with 100mM KCl, 2mM magnesium chloride (MgCl_2_), and 2mM ATP for at least 4 hours at 4°C, and used for further procedures.

### ULP1 peptidase production

Plasmid containing ULP1 peptidase, pFGET19_Ulp1, was a gift from Hideo Iwai (Addgene plasmid # 64697).^61^ After transformation into *E. coli* BL21 (NEB # C2540), cells were grown in terrific broth (Fisher Scientific, Waltham, MA, #BP2468, 50.8 g/L) supplemented with 8mL/L glycerol and 50μg/mL kanamycin to OD600 0.8, and Isopropyl β-D-1-thiogalactopyranoside (IPTG, Fisher Scientific, #15529019) added to 1mM to induce expression. After overnight expression, cells were spun down with centrifugation (4,000xG 10 min.) and pellets frozen at - 80°C, then resuspended in lysis buffer to OD 30. Lysis buffer contained 50mM Tris-hydrochloride (Tris-HCl, pH 8), 300 mM sodium chloride (NaCl), 10mM imidazole, and 1mM TCEP. Cells were lysed using a Branson (Brookfield, CT) SFX 550 sonifier using a 3s on/9s off cycle at 35% amplitude for 10 minutes total. ULP1 was purified using 3mL HisPur Ni-NTA spin columns (Fisher Scientific, #88226) as previously carried out,^62^ and elution fractions were stored in 50% glycerol at -20°C.

### Myosin protein production

Plasmids containing sequence for bovine myosin fragments were transformed into *E. coli* BL21 (NEB # C2540) and transformed bacteria were kept on kanamycin-containing agar-LB broth plates at 4°C or in 30% glycerol (Sigma Aldrich G6279) for long-term storage at -80°C. Terrific broth (Fisher Scientific, Waltham, MA, #BP2468, 50.8 g/L), supplemented with 8mL/L glycerol and 50μg/mL kanamycin, was inoculated with overnight culture to optical density 600nm (OD600) of 0.1, grown to OD600 0.8, and Isopropyl β-D-1-thiogalactopyranoside (IPTG, Fisher Scientific, #15529019) added to 1mM to induce expression. After overnight expression, cells were spun down with centrifugation (4,000xG 10 min.) and pellets frozen at -80°C, then resuspended in lysis buffer to OD 30. Lysis buffer contained 50mM Tris-hydrochloride (Tris-HCl, pH 7.4), 1mM ethylenediaminetetraacetic acid (EDTA), 0.6M KCl, and 0.5mM dithiothreitol (DTT). Cells were lysed using a Branson (Brookfield, CT) SFX 550 sonifier using a 3s on/9s off cycle at 25% amplitude for 12 minutes total. Protein was precipitated using either 6-fold dilution with de-ionized water or dialysis against dialysis buffer using SnakeSkin dialysis tubing with 10k kDa cut-off (Fisher Scientific #88245). Dialysis buffer contained 10mM sodium phosphate (pH 7.4), 0.1M KCl, and 0.5mM DTT. Precipitated protein was spun down (3,700xG 20 min.) and used for further procedures. For freeze-drying, samples were lyophilized in LabConco (Kansas City, MO) Freezone for 48 hours.

### Protein concentration determination

Protein concentration was determined using standard protocols with either Bio-Rad (Hercules, CA) protein assay dye reagent (#5000006) or Pierce BCA protein kit (Fisher Scientific #23225) based on buffer compatibility, with bovine serum albumin standards.

### SDS-PAGE Analysis

Protein samples were boiled in reducing Laemmli SDS buffer (Fisher Scientific, #J61337.AC) for 10 minutes and run on Novex Tris-Glycine 4-20% gradient protein gel (Fisher Scientific #XP04205) at 130V for 90 minutes. Gels were rinsed and stained using SimplyBlue SafeStain (Fisher Scientific #LC6065) according to manufacturer protocols, and imaged using Bio-Rad ChemiDoc Imaging System. Relative protein purity was assessed using band densitometry with ImageJ/Fiji software version 1.53 Gel Analyzer tool.

### Phalloidin staining

F-actin was stained with AlexaFluor Phalloidin 488 (Fisher Scientific #A12379) at 3.3μM concentration. Staining was carried out for 1 hour at room temperature and stained filaments were imaged using a glass coverslip with BZ-X810 (KEYENCE, Osaka, Japan).

### Tube inversion assay

To test heat-induced gel formation ability of expressed myosin fragments, we used established tube inversion assay methods.^63^ A small sample of protein was placed in a 2mL Eppendorf (Hamburg, Germany) tube and heated to 90°C, then allowed to set overnight at 4°C. Tubes were inverted and photos captured using iPhone 16 (Apple, Cupertino, CA).

### Fiber microscopy and analysis

Myosin and actin fibers were using brightfield microscopy using BZ-X810 microscope. (KEYENCE, Osaka, Japan). Actin fibers were also imaged using scanning electron microscopy (SEM, Ultra 55 field-emission scanning electron microscope, Zeiss Gemini 560 FE-SEM, Jena Germany) using 10kV voltage after drying in desiccation chamber and sputter coating with 5nm thing layer of Pt/Pd. Fiber diameter was determined using ImageJ/Fiji software version 1.53 measurement tool.

### Texture profile analysis and sample preparation

Soy protein isolate (SPI) was purchased from Bulk Supplements (Henderson, NV). Myofibrillar proteins, SPI, and buffers were mixed manually in 2mL Eppendorf (Hamburg, Germany) tubes to create formulations of 15% by weight solids. Myofibrillar proteins were added to appropriate concentrations. Mixtures were packed by centrifugation at 300xG for 5 minutes in swing-bucket rotors, then heated on a heat block at 90°C for 1 hour before cooling on ice. Cooked samples were removed from tubes and divided into cylindrical samples 8mm in diameter and 6mm in height for texture profile analysis. The full process for sample preparation is in Figure S2. Sirloin beef was purchased from Bob’s Italian Foods (Medford, MA) and samples prepared in the same fashion.

Texture profile analysis (TPA) was carried out using TA.XTPlus 100 connect (Texture Technologies, Hamilton, MA) with 5kg load cell. TPA measurement was carried out using a double-compression test as previously described,^20^ with 3mm/s test speed, testing to 50% strain deformation, with 2s pause between compressions. Measurements for texture (hardness, chewiness, springiness, elasticity, cohesiveness) were determined using Exponent Connect software. Young’s modulus was determined by calculating the slope of force (kPa) versus strain % in the linear region.

### iBSC cultivation and analysis

To determine myofibrillar protein output of cultivated meat, we grew and differentiated immortalized bovine satellite cells (iBSCs) and analyzed whole-cell protein content. iBSCs were previously isolated from the semitendinosus of a 5-week-old Simmental calf at the Tufts Cummings School of Veterinary Medicine, and engineered to constitutively express bovine telomerase reverse transcriptase (TERT) and cyclin-dependent kinase 4 (CDK4) ^64^. Cells were cultured as previously described.^65^ Briefly, cells were passaged in media containing DMEM + Glutamax (Fisher Scientific #10566024) with 20 % FBS (Fisher Scientific #26140079), 1 ng/mL human fibroblast growth factor 2 (FGF-2) (PeproTech #100-18B, Rocky Hill, NJ, USA), supplemented with 1 % antibiotic/antimycotic (Fisher Scientific #1540062). iBSCs were cultured at 37 °C and 5 % humidity on tissue culture polystyrene (TCPS). iBSCs were grown to confluence, the differentiated using DMEM/F12 (Fisher Scientific #11320033) supplemented with 2 % horse serum (ThermoFisher #16050130), 1 × ITS-X (Fisher Scientific #51400056) and 1 % antibiotic/antimycotic. Differentiation continued for four days before cells reached maximum myotube formation. Immunostaining myosin heavy chain and nuclei of differentiated cells was carried out as described,^65^ using goat anti-mouse secondary antibody AlexaFluor 488 (Fisher Scientific #A-11001) 1:100. Whole-cell protein was extracted from differentiated cells using RIPA lysis buffer (Fisher Scientific #89901) supplemented with 1.5% sodium dodecyl sulfate (SDS) and 50mM DTT, according to manufacturer protocols. Soluble and insoluble protein were separated by centrifuging at 16,000xG for 20 minutes. Soluble fraction protein was collected, concentration was measured with Pierce BCA, and protein components were determined via SDS-PAGE. Myosin and actin relative concentration was determined using band densitometry with ImageJ/Fiji software version 1.53 Gel Analyzer tool.

### Data/statistics

Texture analysis data was analyzed using GraphPad Pris version 10.6. Significance measurements were carried out using one-way ANOVA with Dunnett’s correction for multiple comparisons to detect the presence of significantly different groups.

## CONCLUSION

This study demonstrates the use of precision fermentation from microbial hosts to produce actin and myosin, the myofibrillar proteins largely responsible for the taste and texture of conventional meat. We demonstrate improved soluble yield of recombinant actin using a SUMO tag and show improved recovery through precipitation with ammonium sulfate. We also pinpoint a 99-kDa fragment of the myosin tail region with desirable expression and gelling properties.

Recombinant myosin is shown to be easily purified from the supernatant with dilution or buffer exchange. Thus, myofibrillar protein fermentation can be carried out in a scalable manner for food industry applications. The recombinant actin and myosin mimic the morphological and hierarchal structure of meat tissue, important for creating realistic meat-like mouthfeel. These proteins also improve important textural properties of plant-based proteins, improving hardness, cohesiveness, chewiness, and stiffness at just 1% composition, showing their potential use as meat-mimetic food ingredients. Recombinant myofibrillar proteins pose a novel approach to generating functional animal proteins using cells instead of traditional livestock, in a way that is cheaper and more efficient than cultivated meat.

## SUPPORTING INFORMATION

The following files are available free of charge: (See Supporting information for publication)

- SDS-PAGE of neat actin precipitation; sample preparation scheme for SPI texture analysis; SDS-PAGE of iBSC protein; cost breakdown of protein production; primers used for plasmid constructs; codon sequence and diagram for plasmid constructs

### CRediT authorship contribution statement

**James Dolgin** - Conceptualization, formal analysis, funding acquisition, investigation, methodology, supervision, writing – original draft. **Cornelia H. Barrett** - Conceptualization, investigation, methodology, writing – review and editing. **Maxwell J. Nakatsuji** - Investigation, methodology, writing – review and editing. **Juan Aguilera-Moreno** - Methodology, resources, writing - review and editing. **David L. Kaplan** - Conceptualization, funding acquisition, project administration, supervision, writing – review and editing

## Acknowledgements

We thank the USDA (2021-05678) and the DoD NDSEG Fellowship Program for support of this research. We also thank Drs. Constancio Gonzalez-Obeso, Ryan Scheel, and Emily J. Hartzell for their avice and input regarding molecular biology and protein preparation.

## Data availability

Data will be made available on request.

## Table of Contents Graphic

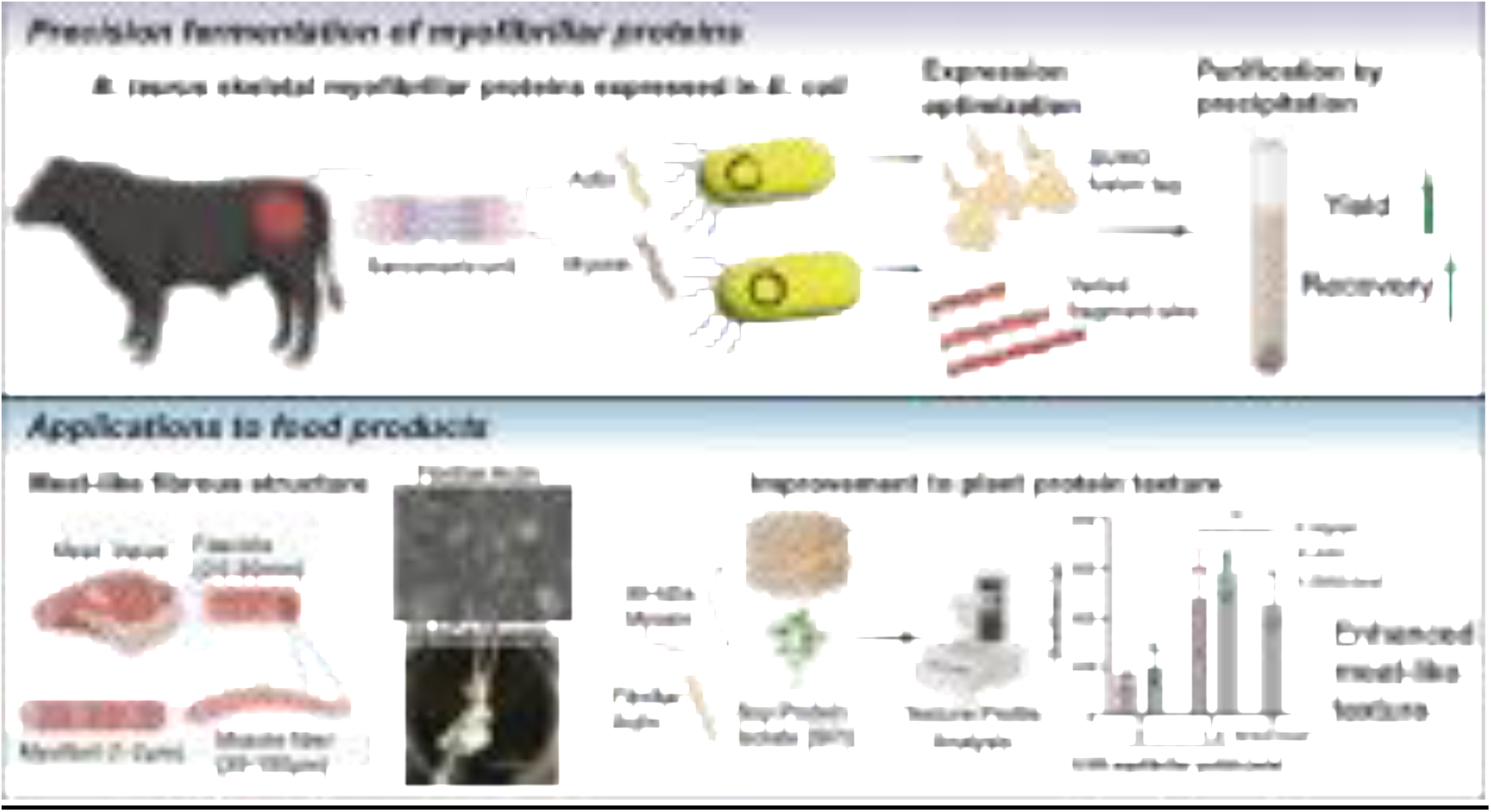

