## Supplemental data for "Precision Fermentation of Recombinant Myofibrillar Proteins for Future Foods"

8 pages; 4 figures; 4 tables

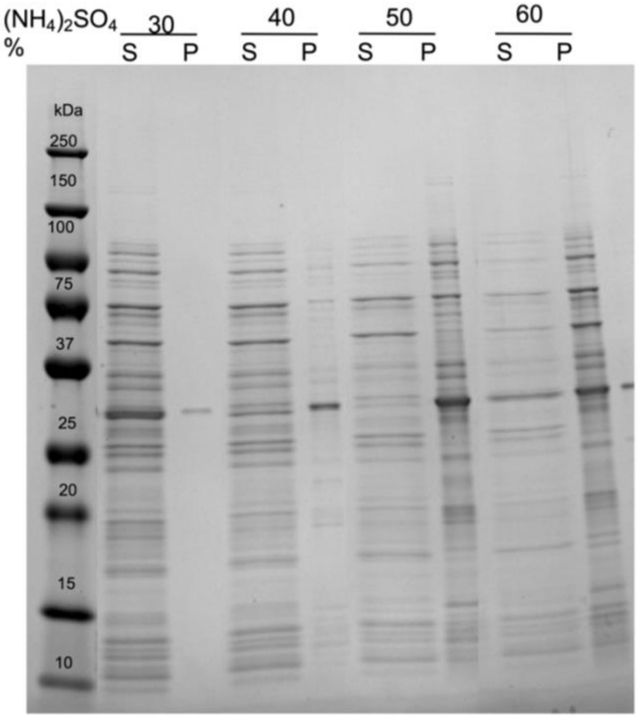


**Figure S1:** SDS-PAGE showing precipitation of non-SUMOylated actin from crude supernatant, showing various concentrations of saturated ammonium sulfate used and respective supernatant (S) and pellet (P) after precipitation and centrifugation.

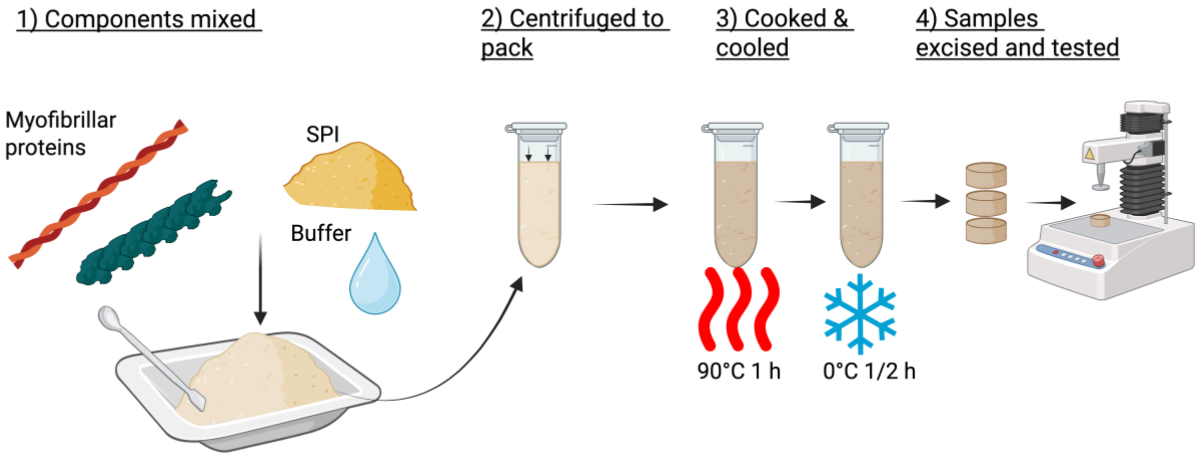


**Figure S2:** Process for sample preparation of SPI & myofibrillar proteins for textural analysis (created with biorender.com).

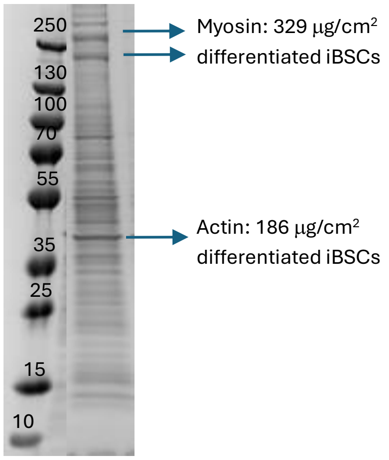


**Figure S3:** SDS-PAGE of whole-cell protein extracted from differentiated immortalized bovine satellite cells (iBSCs) showing yield of myofibrillar proteins normalized to surface area of adherent growth.

| Production stage | Ingredient | Purpose | Supplier | Catalog # | Unit | Cost ($)/unit | Unit needed for 1g myosin | Cost of unit for 1g myosin ($) | Total Cost ($) |
|---|---|---|---|---|---|---|---|---|---|
| Expression | Terrific broth | Carbon source, nutrients | FisherScientific | AAH2682436 | g | 0.282 | 88.81 | 25.04 | |
| | Glycerol | Additional carbon source | FisherScientific | AAA16205AP | ml | 0.11 | 6.99 | 0.77 | |
| | Kanamycin | Selection marker | FisherScientific | 11815032 | g | 7.182 | 0.00 | 0.00 | |
| | IPTG | Expression induction | Sigma Aldrich | I6758 | g | 64.6 | 0.42 | 26.91 | 52.73 |
| Lysis | Tris HCl | Buffer | Sigma Aldrich | 648317 | g | 0.438 | 2.76 | 1.21 | |
| | KCl | Ionic strength | Sigma Aldrich | 529552 | g | 0.114 | 15.64 | 1.78 | |
| | DTT | Reducing agent | Sigma Aldrich | DTT-RO | g | 23.92 | 0.03 | 0.65 | |
| | EDTA | Chelator | Sigma Aldrich | 798681 | g | 0.139 | 0.10 | 0.01 | 3.65 |
| | | | | | | | | **Grand total ($)** | 56.38 |

**Table S1:** Breakdown for estimated raw materials cost needed for 1g myosin production via cell cultivation of iBSCs.

| Production stage | Ingredient | Purpose | Supplier | Catalog # | Unit | Cost ($)/unit | Unit needed for 1g myosin | Cost of unit for 1g myosin ($) | Total Cost ($) |
|---|---|---|---|---|---|---|---|---|---|
| Proliferation | DMEM+glutamax | Basal Media | ThermoFisher | 10566016 | L | 60.84 | 0.96 | 58.51 | |
| | FBS | Growth proteins, peptides, etc. | ThermoFisher | 26140079 | L | 1676 | 0.24 | 408.07 | |
| | Anti-Anti | Antibiotic | ThermoFisher | 15240062 | ml | 0.5465 | 12.17 | 6.65 | |
| | FGF | Growth factor | Peprotech | 100-18B | ug | 0.932 | 1.22 | 1.13 | 474.37 |
| Differentiation | DMEM/F12 | Basal Media | ThermoFisher | 11320033 | L | 67.72 | 1.17 | 79.14 | |
| | Horse serum | Serum for differentiation | ThermoFisher | 16050114 | L | 132.65 | 0.02 | 3.23 | |
| | ITS-X | Differentiation factors | ThermoFisher | 51500056 | ml | 10.4 | 12.17 | 126.61 | 208.98 |
| | | | | | | | | Grand total ($) | 683.35 |

**Table S2:** Breakdown for estimated raw materials cost needed for 1g myosin production via precision fermentation.

| Purpose | PCR Target for Assembly | Fwd. primer (*overhangs in italics)* | Rev. primer (*overhangs in italics)* |
|---|---|---|---|
| Assembly of Actin into pCIOX SUMO plasmid | Vector | 5' *ACAGAGAACAGATTGGTGGT* ATGTGCGACGAAGATGAAAC 3' | 3' *GCTTTGTTAGCAGCCGGATCTTA* AAAACATTTACGATGAACAATACTCGG 5' |
| Assembly of Actin into pCIOX SUMO plasmid | Backbone | 5' GATCCGGCTGCTAACAAAGC 3' | 3' CGAGTGTCTCTTGTCTAACCACCA 5' |
| Removal of SUMO sequence | | 5' ATGTGCGACGAAGATGAAACCAC 3' | 3' GCTAGCGCTGCCGCG 5' |
| Assembly of Myosin 64 kDa into pCIOX plasmid | Vector | 5' *CAGCGCTAGC* TCTAGACTGGAAGAGGCC 3' | 5' GTTTTAATAGTCGCTTCTTGAGCTC *ATTGCGCCTA* 3' |
| Assembly of Myosin 64 kDa into pCIOX plasmid | Backbone | 5' *AGAACTCGAG* TAACGCGGATCCGAATTCGAGCTCCG 3' | 5' GGCGCGCCGTCGCGATCG *AGATCTGACC* 3' |
| Assembly to express 131 kDa myosin | Vector | 5' *GGGCAGCAGC*CGTGAAAGCATTTTCTGC 3' | 3' *CCTCTTCCAG* TTCTTCGGTACGCTGAATTG 5' |
| Assembly to express 131 kDa myosin | Backbone | 5'*TACCGAAGAA* CTGGAAGAGGCCAAAAAGAAACTGGCACAGC 3' | 3' GGCGCGCCGTCG *CGATCGGCACTTTCGT* 5' |
| Shortening myosin to 99 kDa | | 5' ATGAGCAACCTGCAG 3' | 3' GCTGCTGCCCATATG 5' |

**Table S3:** Primers used to construct plasmids in this study.

| | |
|---|---|
| Bovine actin codon sequence | ATGTGCGACGAAGATGAAACCACCGCACTGGTTTGTGATAATGGTAGCGGTCTGGTTAAAGCAGGTTTTG<br>CCGGTGATGATGCACCGCGTGCAGTTTTTCCGAGCATTGTTGGTCGTCCGCGTCATCAGGGTGTTATGGT<br>TGGTATGGGTCAGAAAGATAGCTATGTTGGTGATGAAGCACAGAGCAAACGTGGTATTCTGACCCTGAAAT<br>ATCCGATTGAACATGGCATTATTACCAACTGGGATGACATGGAAAAGATCTGGCATCACACCTTTTATAACG<br>AACTGCGTGTTGCACCGGAAGAACATCCGACACTGCTGACCGAAGCACCGCTGAATCCGAAAGCAAATC<br>GTGAGAAAATGACCCAGATTATGTTCGAAACCTTTAACGTTCCGG<br>CAATGTATGTTGCAATTCAGGCAGTTCTGAGCCTGTATGCAAGCGGTCGTACCACCGGTATTGTTCTGGAT<br>AGCGGTGATGGTGTTACCCATAATGTTCCGATTTATGAAGGTTATGCACTGCCGCATGCAATTATGCGTCTG<br>GATCTGGCAGGTCGTG<br>ATCTGACCGATTATCTGATGAAAATTCTGACCGAACGCGGTTATAGCTTTGTTACCACCGCAGAACGTGAA<br>ATTGTGCGCGATATTAAAGAAAAACTGTGCTATGTTGCCCTGGATTTTGAAAATGAAATGGCAACCGCAGC<br>AAGCAGCAGCAGCCTGGAAAAGAGCTATGAACTGCCGGATGGTCAGGTTATTACCATTGGCAATGAACGT<br>TTTCGTTGTCCGGAAACACTGTTTCAGCCGAGCTTTATTGGTATGGAAAGCGCAGGTATTCATGAAACGAC<br>CTATAACAGCATCATGAAATGCGATATTGACATCCGCAAAGATCTGTATGCCAATAATGTTATGAGCGGTGG<br>CACCACCATGTATCCGGGTATTGCAGATCGTATGCAGAAAGAAATTACAGCACTGGCACCGAGCACCATGA<br>AAATCAAAATCATTGCACCGCCTGAACGCAAATATAGCGTTTGGATTGGTGGTAGCATTCTGCGTAGCCTG<br>AGCACCTTTCAGCAGATGTGGATTACCAAACAAGAATATGATGAAGCCGGTCCGAGTATTGTTCATCGTAA<br>ATGTTTTTAA |
| Bovine myosin (residues 811-1938) codon sequence | CGTGAAAGCATTTTCTGCATTCAGTATAATGTGCGTGCCTTCATGAATGTTAAACATTGGCCGTGGATGAAG<br>CTGTACTTTAAGATTAAACCGCTGCTGAAATCAGCCGAAACCGAAAAAGAAATGGCCAACATGAAAGAAGA<br>GTTTGAGAAAACCAAAGAGGAATTGGCCAAAAGCGAAGCCAAACGCAAAGAACTGGAAGAAAAGATGGT<br>TACCCTGACGCAAGAGAAAAATGATCTGCAGTTACAGGTTCAGAGCGAAGCAGATGCACTGGCCGATGCC<br>GAAGAACGTTGTGACCAGCTGATCAAAACCAAAATTCAACTGGAAGCCAAGATCAAAGAAGTTACCGAAC<br>GCGCAGAAGATGAAGAAGAAATTAACGCAGAACTGACCGCCAAAAAGCGTAAGCTGGAAGATGAGTGTAG<br>CGAACTGAAAAAGGATATCGATGATCTGGAACTGACCCTGGCCAAAGTTGAGAAAGAAAAACATGCCACC<br>GAGAACAAAGTGAAAAATCTGACCGAAGAGATGGCAGGTCTGGATGAAACCATTGCCAAACTGACCAAAG<br>AAAAGAAAGCACTGCAAGAAGCCCATCAGCAGACCCTGGATGACCTGCAGGCAGAAGAAGATAAAGTTAA<br>TACCCTGACAAAAGCCAAGACGAAACTGGAACAACAGGTTGATGACCTGGAAGGTTCATTAGAACAAGAG<br>AAGAAACTGCGCATGGACCTGGAACGTGCAAAACGCAAACTGGAAGGTGATCTGAAACTGGCCCAAGAA<br>AGTACCATGGATATCGAAAATGATAAGCAGCAACTGGATGAGAAGCTTAAAAAGAAAGAATTTGAAATGAGC<br>AACCTGCAGAGCAAAATTGAAGATGAACAGGCCCTGGCAATGCAACTGCAGAAAAAGATTAAAGAGCTGC<br>AGGCACGCATTGAAGAACTTGAGGAAGAAATTGAAGCGGAACGTGCCAGCCGTGCAAAAGCCGAAAAAC<br>AGCGTAGCGATCTGAGTCGTGAACTTGAAGAAATTTCAGAACGTTTAGAAGAAGCAGGCGGAGCAACCAG<br>CGCACAGATCGAAATGAATAAGAAACGTGAAGCGGAGTTCCAGAAGATGCGTCGCGATTTAGAAGAGGCA<br>ACCCTGCAGCATGAAGCAACCGCAGCAGCACTGCGTAAGAAACATGCCGATAGCGTTGCAGAACTGGGT<br>GAGCAGATTGATAACCTGCAGCGTGTGAAACAAAAACTGGAAAAAGAAAAATCCGAAATGAAAATGGAAAT<br>CGACGACCTGGCAAGCAATATGGAAACCGTTAGCAAAGCCAAAGGTAACCTGGAAAAGATGTGTCGTGCG<br>CTTGAAGATCAGCTGAGTGAGCTGAAAACAAAAGAAGATGAGCAGCAGCGTCTGATTAATGATCTGACCA<br>CACAGCGTGCTCGCCTGCAGACCGAATCAGGTGAATTTAGCCGTCAGCTGGACGAAAAAGATGCCCTGG<br>TTAGCCAGCTGAGCCGTGGTAAACAGGCATTTACCCAGCAAATAGAAGAACTGAAACGTCAGCTTGAAGA<br>AGAGATCAAAGCAAAAAGCGCACTGGCACATGCGCTGCAGAGCGCACGTCATGATTGTGATCTGCTGCG<br>TGAACAGTATGAAGAGGAACAAGAGGGTAAAGCCGAACTGCAACGTGCAATGAGCAAAGCGAATAGCGA<br>AGTTGCCCAGTGGCGTACCAAATATGAAACCGATGCAATTCAGCGTACCGAAGAACTGGAAGAGGCCAAA<br>AAGAAACTGGCACAGCGTCTGCAGGATGCCGAAGAACATGTTGAAGCAGTTAATGCAAAATGTGCAAGCC<br>TGGAAAAGACCAAACAGCGCCTGCAGAATGAAGTTGAAGATCTGATGATTGATGTGGAACGTACCAATGC<br>AGCATGTGCAGCACTGGATAAAAAGCAGCGTAACTTTGATAAAATCCTGAGCGAGTGGAAGCAGAAATATG<br>AAGAAACCCATGCAGAACTGGAAGCAAGCCAGAAAGAATCACGTAGCCTGAGCACCGAACTGTTCAAAAT<br>CAAAAATGCATATGAAGAGAGCCTGGATCAGCTGGAAACCCTGAAACGTGAAAACAAAAATCTGCAGCAA<br>GAAATCAGCGATCTGACCGAGCAGATTGCAGAAGGTGGTAAACGTATTCATGAACTGGAAAAAGTGAAGA<br>AACAGGTGGAACAAGAGAAAAGCGAAATTCAGGCAGCCCTGGAAGAAGCCGAAGCCAGCCTGGAACAT<br>GAAGAAGGTAAAATTCTGCGTATTCAGCTTGAACTGAATCAGGTGAAAAGTGAGATTGATCGCAAAATCGC<br>CGAAAAAGATGAGGAAATTGATCAGCTGAAGCGCAACCATATTCGTATTGTTGAAAGCATGCAGAGCACCC<br>TGGATGCAGAAATTCGTAGCCGTAATGATGCAATTCGGCTGAAAAAGAAAATGGAAGGCGATCTGAACGAA<br>ATGGAAATCCAGCTGAATCATGCAAATCGTATGGCAGCAGAAGCACTGAAAAACTATCGTAGCACCCAGGC<br>AATTCTGAAAGATACCCAGATTCATCTGGATGATGCACTGCGTGGTCAAGAGGATCTGAAAGAACAGCTG<br>GCAATGGTTGAACGTCGTGCAAATCTGCTGCAGGCCGAAATTGAAGAACTGCGTGCGACCCTGGAACAG<br>ACCGAACGTAGTCGTAAAATTGCAGAACAAGAACTGCTGGATGCAAGCGAACGTGTTCAGCTGCTGCATA<br>CCCAGAATACCAGCCTGATTAATACCAAGAAAAAGCTTGAAACCGACATCACACAGATTCAGGGTGAAATG<br>GAAGATATTATTCAAGAAGCACGTAACGCAGAAGAGAAAGCCAAAAAGGCAATTACCGATGCAGCAATGAT<br>GGCGGAAGAACTGAAGAAAGAGCAGGATACCAGCGCACATCTGGAACGCATGAAGAAAAATCTGGAACA<br>AACCGTTAAAGACCTGCAGCATCGTCTGGATGAAGCAGAGCAGCTGGCACTGAAAGGTGGCAAAAAGCA<br>GATCCAGAAATTAGAAGCCCGTGTGCGTGAGCTGGAAGGTGAAGTGGAAAGCGAACAGAAACGTAATGT<br>GGAAGCCGTTAAAGGTCTGCGTAAACATGAACGTCGCGTTAAAGAACTGACCTATCAGACCGAAGAAGAT<br>CGGAAAAACATTCTGCGTCTGCAAGATCTGGTTGATAAACTGCAGGCAAAAGTGAAAAGCTATAAACGTCA<br>GGCAGAAGAAGCAGAGGAACAGAGCAATGTTAATCTGAGCAAATTTCGCAAACTGCAGCATGAATTAGAA<br>GAGGCTGAAGAACGTGCAGATATTGCAGAAAGCCAGGTGAATAAACTGCGTGTTAAAAGCCGTGAAGTGC<br>ACACCAAAATTATCAGCGAAGAATAA |

**Table S4:** Optimized codon sequences for bovine actin used in this study, and optimized codon sequences for myosin tail fragment showing 64 kDa fragment (no highlight), added codons for 99 kDa fragment (blue highlight), and added codons for 131 kDa fragment (green highlight)

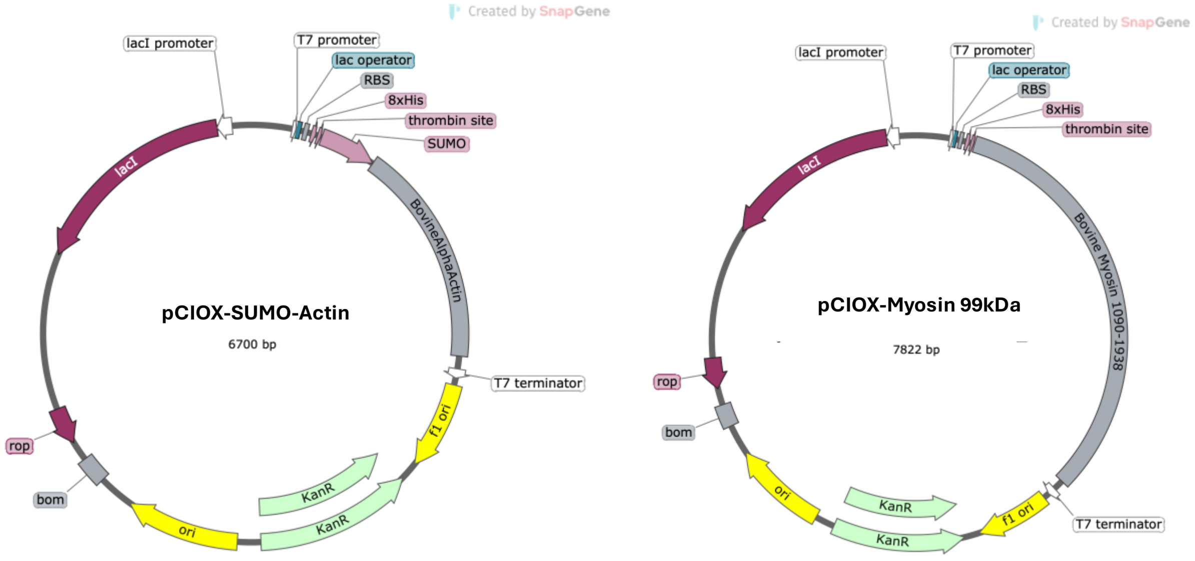


**Figure S4:** Constructs used for expression of SUMO-Actin (left) anf 99-kDa myosin (right)
